# Longitudinal TCR repertoires in ulcerative colitis patients show features distinguishing disease states

**DOI:** 10.1101/2025.07.31.667918

**Authors:** Sabahat Rahman, Matthew Farah, Amanda Kwok, Jacob Varghese, Beiya Xu, Amir Daraie, Joseph Greenstein, Casey Overby Taylor, Nidhi Soley, April Yan, Ishan Vatsaraj, Carl Harris, Joanna Melia, Kristi Briggs

## Abstract

**Background:** Ulcerative colitis (UC) affects an estimated 10 million people worldwide. The exact etiology of the disease is unknown, but T cell dysregulation and aberrant activation is associated with UC—prompting research into T cell receptor (TCR) repertoires in UC patients. However, few studies have compared UC patients’ TCR repertoires in flare (inflamed) and remission (uninflamed) states. Moreover, this is the first dataset to our knowledge examining multiple repertoires from UC patients over time, enabling longitudinal analyses.

**Methods:** TCR repertoires were obtained for 21 patients across multiple timepoints, yielding a total of 58 samples. Repertoires’ clonality, diversity, overlap, and gene usage frequencies were compared across all patients. CDR3 sequences were split into K-mers (sequences of length K), and enriched K-mers in flare and remission states were identified using GLIPH2.

**Results:** Although repertoires vary across patients, there were significant differences in the usage of 20 Vβ genes across flare and remission states. Moreover, calculating the overlap in repertoires with enriched K-mers showed higher overlap scores than when using full TCR sequences. 622 unique enriched K-mers were identified in flare states and 495 in remission states, with only 57 overlapping between the two states.

**Conclusions:** Overall, these results highlight the importance of analyzing Vβ gene usage and K-mer enrichment in TCR repertoires, particularly given the lack of public clones across patient cohorts. Future studies characterizing the specific antigenic targets associated with these features will pave the way for biomarker discovery in UC.

**KEY MESSAGES:**

- **What is already known?** UC is a chronic condition characterized by fluctuating periods of flares and remissions, as well as aberrant immune activity.
- **What is new here?** TCR repertoires were obtained from UC patients’ colonoscopies, with multiple timepoints per patient—enabling the analysis of repertoire characteristics within and across patients over time.
- **How can this study help patient care?** Understanding the exact immune mechanisms in UC can pave the way for biomarker discovery and advance the state of care for patients, given that current treatments are non-specific and have adverse side effects.

**Summary:** This article presents an analysis of T cell receptor repertoires from ulcerative colitis patient samples collected during colonoscopies, identifying genes and motifs associated with patients’ flare (inflamed) and remission (uninflamed) states.

## INTRODUCTION

The two main inflammatory bowel diseases (IBD) are Crohn’s disease (CD) and ulcerative colitis (UC), which are characterized by distinct pathophysiological characteristics and clinical features. CD can involve any segment of the gastrointestinal (GI) tract, with a tendency to cause transmural inflammation of the GI mucosa, including fistula formation. UC, on the other hand, is associated with inflammation mainly in the rectum and colon.^1^ UC is of particular interest because of its rising incidence, especially in the developed world, and its distinctive pattern of colonic involvement, which presents difficulties both in diagnosis and in therapeutic management.^2^ The disease follows a chronic course—patients fluctuate between bouts of active inflammation and temporary remission—which renders disease progression unpredictable and makes treatment difficult. Current treatment relies on a variety of pharmacologic agents, including anti-inflammatory drugs, immunosuppressants, and biologic agents, all aimed at lessening inflammation and inducing remission. Although these drugs have drastically reduced mortality, they are not UC-specific and possess general immunosuppressive effects, with a tendency to have severe adverse effects.^2,3^ In addition, UC disease monitoring predominantly utilizes nonspecific inflammatory markers, such as C-reactive protein and erythrocyte sedimentation rate.^4^ Although helpful for clinicians, these markers do not illuminate the specific immune processes involved in UC and are confounded by their elevation in other inflammatory conditions. Finally, despite the advancements and increases in treatment and disease monitoring, a significant number of patients have persistent symptoms and fail to benefit from these therapies—Le Berre et al. (2023) estimates 10-20% of UC patients must undergo proctocolectomy for their disease.^2^

The lack of targeted therapeutics for UC is complicated by the fact that the exact pathogenesis and etiology remain elusive, though genetic susceptibility, environmental influences, and immune system dysfunction are all thought to play a role.^3^ T cell activity in particular has been widely studied. It is widely accepted that abnormal T cell responses may be an underlying factor in UC.^3^ This is logical, given that increased T cell reactivity against antigens presented on peptide-MHC complexes in the gut would drive excess inflammatory signaling, and presents a promising avenue for studying T cell receptors (TCRs) and designing targeted UC therapeutics. Characterizing disease-associated T cell populations via bulk TCR sequencing has been made possible thanks to the advent of accessible sequencing technologies, and TCR repertoire analysis has been employed to study IBDs, COVID-19, multiple sclerosis, ankylosing spondylitis, and cancer.^5-9^ Although previous studies have examined TCR repertoires in UC, these have largely adopted a cross-sectional approach to TCR repertoire analysis—providing static information about the immune environment in patients.^10,11^ However, such methods are unable to account for the dynamic changes characteristic of UC, whereby patients oscillate between inflamed and remission phases. Consequently, long-term TCR repertoire evolution and correlation with disease activity are poorly characterized. The scale of TCR diversity also emphasizes the need for more personalized analyses of patients’ repertoires, which can be achieved by studying multiple repertoires from individual patients over time.

Here we present novel results from the longitudinal assessment of TCR repertoires in UC patients. We sought to investigate repertoire characteristics associated with patients’ flare and remission states, thereby identifying potential markers associated with each disease state. We also investigated repertoire characteristics in patients over time, revealing key insights about changes in patients’ T cell populations. Further analyses like these may allow for a more complete understanding of immune system dynamics in UC patients, as well as inform the development of personalized biomarkers for disease monitoring and treatment—ultimately paving the way to break the “therapeutic ceiling” in UC.

## MATERIALS AND METHODS

### BIOPSY COLLECTION AND SAMPLE PROCESSING

All formalin-fixed paraffin-embedded (FFPE) samples were obtained from UC patients at the Johns Hopkins Hospital and sequenced under an IRB approved protocol (IRB 00336979). Patients who a) had colonoscopy biopsy specimens from the sigmoid colon and b) whose samples were categorized as coming from periods of active disease or remission were included in this study. Although 23 patients were initially identified, 21 patients’ sequencing data were used for final analyses—one patient did not have a sufficient sequencing depth, and another did not have enough FFPE sample for analysis. Some patient samples collected at specific timepoints were also dropped because of issues with sequencing depth or FFPE availability. Finally, a total of 58 samples from 21 patients were included in analyses, with 2-4 available time points for every patient except Patient 5 (only one time point). Data spanned nine years and were obtained from a diverse patient cohort (**Table 1**).^12^

**Table 1.** Summary of all patients’ metadata.

| Patient Number | Age (years) | Gender | Disease Duration (years) | Time period between the first and last biopsy (years) | Number of biopsies |
| --- | --- | --- | --- | --- | --- |
| 1 | 68 | M | 19 | 6 | 4 |
| 2 | 53 | F | 3 | 1 | 2 |
| 3 | 26 | M | 7 | 3 | 2 |
| 4 | 39 | F | 12 | 2 | 2 |
| 5 | 43 | F | 8 | N/A | 1 |
| 6 | 68 | M | 6 | 3 | 3 |
| 7 | 59 | F | 8 | 2 | 2 |
| 8 | 29 | F | 5 | 1 | 2 |
| 9 | 34 | M | 9 | 5 | 4 |
| 10 | 43 | F | 5 | 1 | 2 |
| 11 | 39 | F | 11 | 5 | 4 |
| 12 | 41 | F | 22 | 0 yr 7 mon | 2 |
| 13 | 69 | F | 26 | 9 | 3 |
| 14 | 71 | F | 19 | 2 | 3 |
| 15 | 37 | F | 26 | 4 | 3 |
| 16 | 35 | M | 8 | 3 yr 5 mon | 3 |
| 17 | 25 | M | 6 | 1 | 2 |
| 18 | 37 | M | 7 | 6 | 3 |
| 19 | 24 | M | 10 | 5 | 4 |
| 20 | 50 | F | 26 | 5 | 3 |
| 21 | 40 | M | 9 | 4 | 4 |
| Summary (Mean ± SD) | 43.7 ± 14.3 | — | 11.0 ± 7.7 | 3.6 ± 2.0 (excluding N/A) | 2.8 ± 0.9 |

Samples were classified as flares or remissions based on the following criteria. Flares were defined as pathologically active disease states with Mayo Endoscopic Scores (MES) of 2 or 3 from endoscopic analysis, whereas remissions were defined as inactive disease states with MES of 0 or normal colonic mucosa.^13^ Differences in pathological and endoscopic findings were rare but addressed by reviewing cases with a pathologist to determine a final classification.^12^

### TCR SEQUENCING AND REPERTOIRE ANALYSIS

DNA was extracted from FFPE-preserved sigmoid colon biopsy samples using the Qiagen DNeasy Blood and Tissue Kit. TCRβ chains were PCR-amplified and sequenced by the Johns Hopkins Sidney Kimmel Comprehensive Cancer Center FEST and TCR Immunogenomics Core Facility using the Ampliseq for Illumina TCR-short read kit. All sequencing and data quality control was done as previously described.^12,14^

All sequencing data were converted to VDJtools format for subsequent analyses. Samples are labeled and referred to as PtX_[F/R]_[1/2/3/4], where X denotes the patient ID, F/R represent flare or remission states, respectively, and the final number represents the biopsy’s ordinal position. Repertoire clonality, diversity, and overlap, as well as all alluvial plots, were calculated and visualized using Immunarch (v0.9.1). Ambiguous gene assignments (ex. TRBV12-3, TRBV12-4) were represented with their broader family name (ex. TRBV12 in the former case) for further analyses. To investigate differential gene usage patterns, the mean frequency of each gene across all flare states and all remission states was calculated (**Eq. 1**). Statistical comparison for the frequencies of each gene family between flare and remission groups was performed using the Mann–Whitney U test; multiple testing was corrected using the Benjamini–Hochberg procedure. All analyses were done in R (v4.4.1) and Python (v3.12.10).

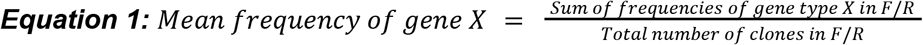

### GLIPH MOTIF ANALYSIS

The turboGLIPH R package, an optimized version of the GLIPH algorithm originally developed by Glanville et al. (2017), was used to identify enriched K-mers in individual patient repertoires using the default parameters of the algorithm.^15,16^ HLA data were not included. A key output of GLIPH is its “motif_enrichment” output. This is a list of two data frames containing selected (enriched) motifs and all motifs found in the input repertoires. To minimize an inflated overlap in K-mers between different patients, only the selected motifs—which passed the GLIPH p-value and observed versus expected ratio criterion defined by turboGLIPH’s gliph2 function—were used for overlap analysis. Further, to account for double-counting a single patient’s enriched K-mers, the set of all enriched K-mers found across all TCR repertoires belonging to a single patient was recorded.

Jaccard overlap was employed to measure the similarity between patients based on their enriched motifs. While the Jaccard score does not account for the frequency of each motif, it provides an interpretable measure of the extent to which unique k-mers are shared across different patients.

### DATA AVAILABILITY

All pre-processed and de-identified TCR repertoires will be made available for public access upon publication.

## RESULTS

### SUMMARY STATISTICS DO NOT DIFFERENTIATE FLARE AND REMISSION REPERTOIRES

Metadata for all 21 patients are summarized in **Table 1**. To characterize patients’ TCR repertoires, the number of unique T cell clones (defined by their unique CDR3 sequences) and clonality of each repertoire were calculated (**Figure 1a,b**). Repertoire diversity was calculated using the Chao1 score, which quantifies species richness by accounting for the total number of observed clones, number of singletons (clones occurring once), and doubletons (clones occurring twice) (**Figure 2a**).^17^ Results demonstrated that there were notable intra-patient trends—for example, all of Patient 1’s samples had low Chao1 scores from 187 to 463, demonstrating that they consistently had few clones present at higher frequencies. Indeed, the Pt1_F_1 repertoire has seven clones present at frequencies greater than 0.02—in stark contrast to a repertoire like Pt17_F_1, which had a significantly higher Chao1 score of 2653 and no clones with frequencies greater than 0.0108. Overall, the mean Chao1 score was 1749, and the standard deviation was 1652, further underscoring the high variability across samples. These findings suggest that T cell activity in some UC patients may be driven by expanded clones responding to target antigens, whereas other patients may have less clonal expansion and a more diverse set of antigens in the colon.^18,19^ Nonetheless, across all samples, there were no significant differences in diversity scores across flare and remission states (**Figure 2b**), so differences in T cell CDR3 sequence frequency/expansion are not necessarily associated with inflamed tissue states in this patient cohort.

**Figure 1.**
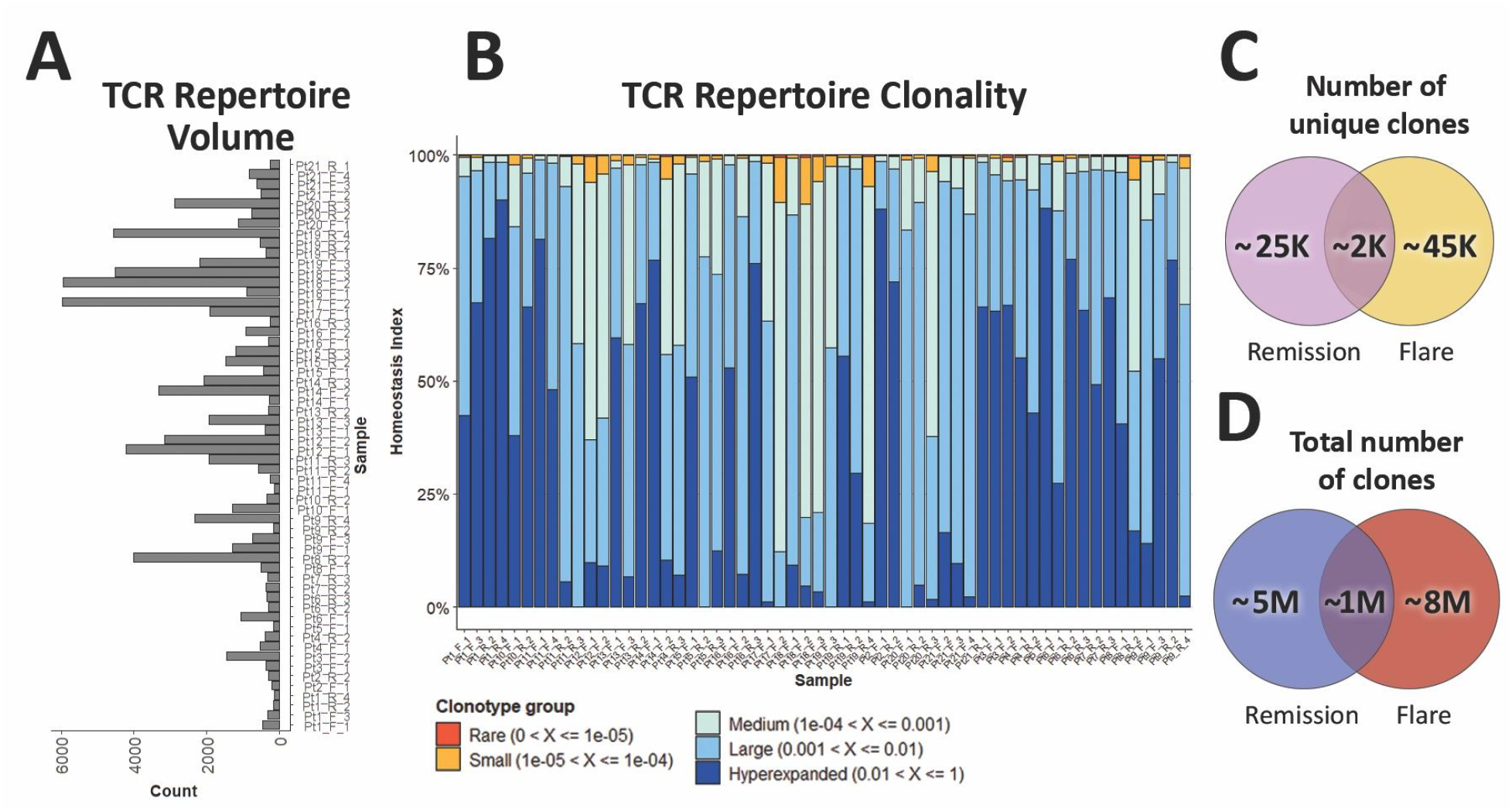
Summary statistics describing all 58 samples. (A) TCR repertoire volume and (B) relative abundance of clonotypes are shown for all 58 samples. (C) The number of unique clones in flare and remission states, as well as (D) the total number of clones across all flare/remission repertoires is shown. **Alt text:** Subfigures A-B show bar plots describing TCR repertoires’ number and abundance of clones. Subfigures C-D show Venn diagrams with the number of unique clones and total number of clones.

**Figure 2.**
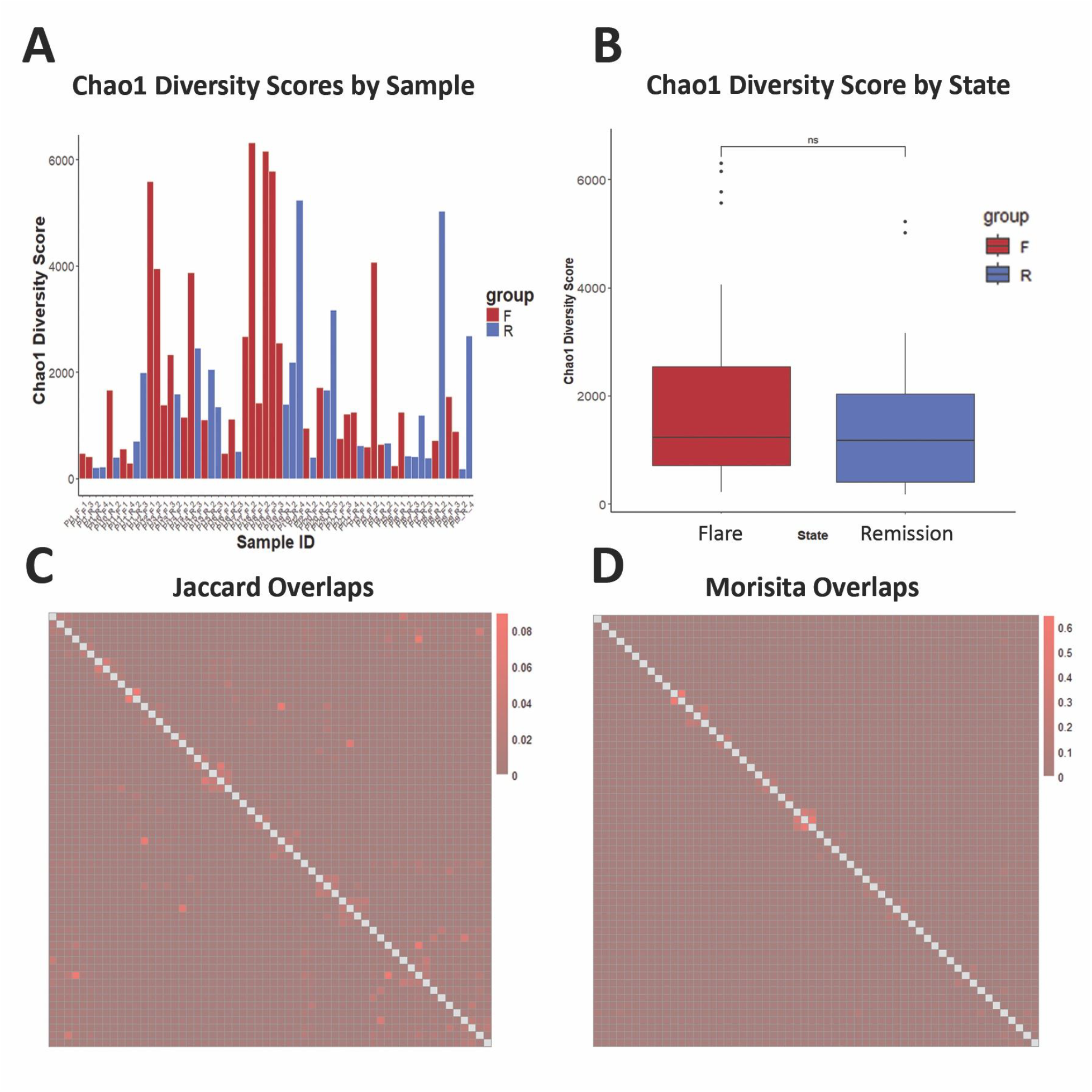
Diversity and overlap scores show substantial heterogeneity across repertoires. (A) Chao1 diversity scores were calculated for all samples. (B) Unpaired two-sided Wilcoxon rank sums test was performed to compare Chao1 scores across all flare (F) and remission (R) samples; ns = non-significant (p>0.05). (C) Jaccard and (D) Morisita overlap scores were calculated across all pairs of samples. **Alt text:** Subfigure A shows a bar plot with Chao1 diversity scores across patients, and B shows a box plot comparing mean Chao1 scores in flare and remission states. C and D show heatmaps with Jaccard and Morisita overlap scores, respectively.

Next, overlap between each pair of TCR repertoires was calculated using the Jaccard and Morisita indices (**Figure 2c,d**). While the Jaccard index is a simple ratio of the size of the intersection and the size of the union of two repertoires, the Morisita index further accounts for the abundance of clones in a repertoire—therefore, higher Morisita indices indicate repertoires are more similar in their composition and abundance.^20^ Little inter-patient similarity was observed, which is reflective of the immense diversity in TCRβ sequences. Slightly higher similarity values were found when comparing samples from individual patients, but the low values overall suggest that patient T cell repertoires change substantially over time and that each disease time point is characterized by its own unique T cell response.

Overall, these initial findings suggest that summary statistics, though interpretable and useful in understanding individual repertoires, are not sufficient for identifying cohort-level trends—particularly trends that may distinguish between flare and remission states and therefore have clinical relevance across patients.

### THERE ARE DIFFERENTIAL GENE USAGE PATTERNS ACROSS FLARE AND REMISSION STATES

Gene usage is an important metric for TCR repertoire analysis, given that individual TCRβ sequences are composed of specific variable (V), diversity (D), and joining (J) segments. The Vβ gene in particular has been widely studied because of its crucial role in TCR recognition of peptide-MHC complexes. Dash et al. (2017) used large-scale TCR sequencing combined with machine learning to show that CDR3 sequences and Vβ gene usage are predictive of TCR antigen specificity.^21^ Similarly, Springer et al. (2021) also showed that CDR3 and Vβ sequences were the features with the greatest contributions in their machine learning model for predicting TCR-peptide binding.^22^

To examine V gene usage patterns in our patient cohort and potential recurring TCR features that distinguish flare and remission states, differential gene usage patterns between the two disease states of UC were assessed (**Figure 3**). The most frequently used Vβ gene families in both flare and remission repertoires were TRBV20 (specifically TRBV20-1), TRBV6, and TRBV7. Several TRBV segments and families, including TRBV28, TRBV20-1, TRBV7, and TRBV27, showed higher mean frequencies during remissions. Overall, significant differences in the utilization of 20 TRBV segments/families, including TRBV28, TRBV6, and TRBV20-1, were found across all flare and remission samples.

**Figure 3.**
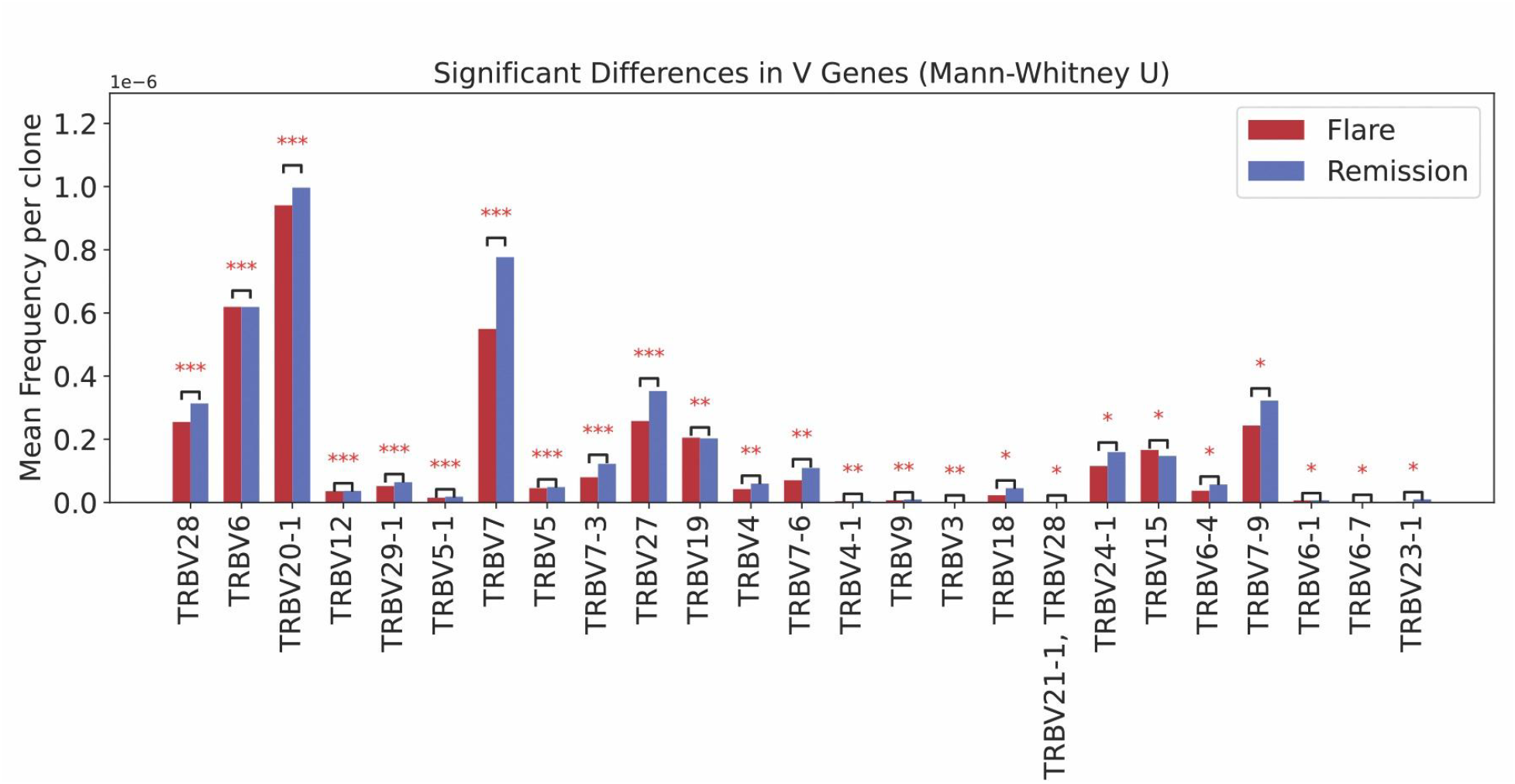
There are differential gene usage patterns across flare and remission states. Results of unpaired two-sided Wilcoxon rank sums test on Vβ gene usage across disease states. Mean frequencies per clone were plotted for all flare and all remission samples. ^*^ = p<0.05, ^**^ = p<0.01, and ^***^ = p<0.001. **Alt text:** Bar plot showing mean frequency of Vβ gene usage across flare and remission samples, with statistical analyses shown on the graph.

### ENRICHED K-MERS ARE UNIQUE TO FLARE AND REMISSION STATES

To further identify characteristic motifs across patients’ TCRs, we analyzed K-mers (sequences of length K) in repertoires’ CDR3 sequences. K-mer analysis is a popular approach in natural language processing and has been extensively applied in omics analysis, particularly with TCR repertoires.^23^ Ostmeyer et al., for instance, demonstrated that 4-mers from TCRβ CDR3 sequences could be used in a machine learning model to classify healthy and cancerous TCR repertoires.^24^ Similarly, Park et al. found K-mer features associated with different COVID-19 disease severities.^6^ Identifying these CDR3 K-mers in specific disease states or patients is of interest, as enriched or shared K-mers may indicate similar antigen specificities across T cell clones.^6,24^ Moreover, K-mer analysis can overcome the hurdle of comparing repertoires when there are few public clones.

Enriched K-mers were identified from patient TCR repertoires using GLIPH2 in two separate analyses. In the first, for each patient, all available repertoires (up to four per individual) were combined, and the GLIPH2 algorithm was run on this set. The resulting enriched K-mers for each patient were then compared pairwise across all other patients. In this analysis, Jaccard similarity scores computed between the unioned repertoires of each patient pair indicated up to 15% overlap in enriched K-mers between some individuals (**Figure 4a**), representing a notable increase in inter-patient similarity compared to full TCR sequence comparisons, where we saw at max 8.9% overlap (**Figure 2c**). To assess whether the enriched K-mers driving this overlap exhibited specificity based on disease activity, GLIPH2 was rerun on patient repertoires stratified by disease activity state (**Figure 4b**). Flare-associated K-mers were compared independently from remission-associated K-mers. The overall Jaccard similarity between the union of all flare K-mers and the union of all remission K-mers was approximately 0.05, amounting to 57 overlapped K-mers out of a set size of 1117 uniquely enriched K-mers—suggesting that K-mer overlaps may reflect distinct immune states rather than random sharing. Specifically, when considering all enriched K-mers, 622 unique K-mers were identified in flares and 495 in remissions, with only 57 overlapping between the two states. This is consistent with results from previous work, wherein K-mers could be used to categorize TCR repertoires as coming from healthy or diseased patients^24^—highlighting their value in capturing distinct immune states rather than reflecting random overlap.

**Figure 4.**
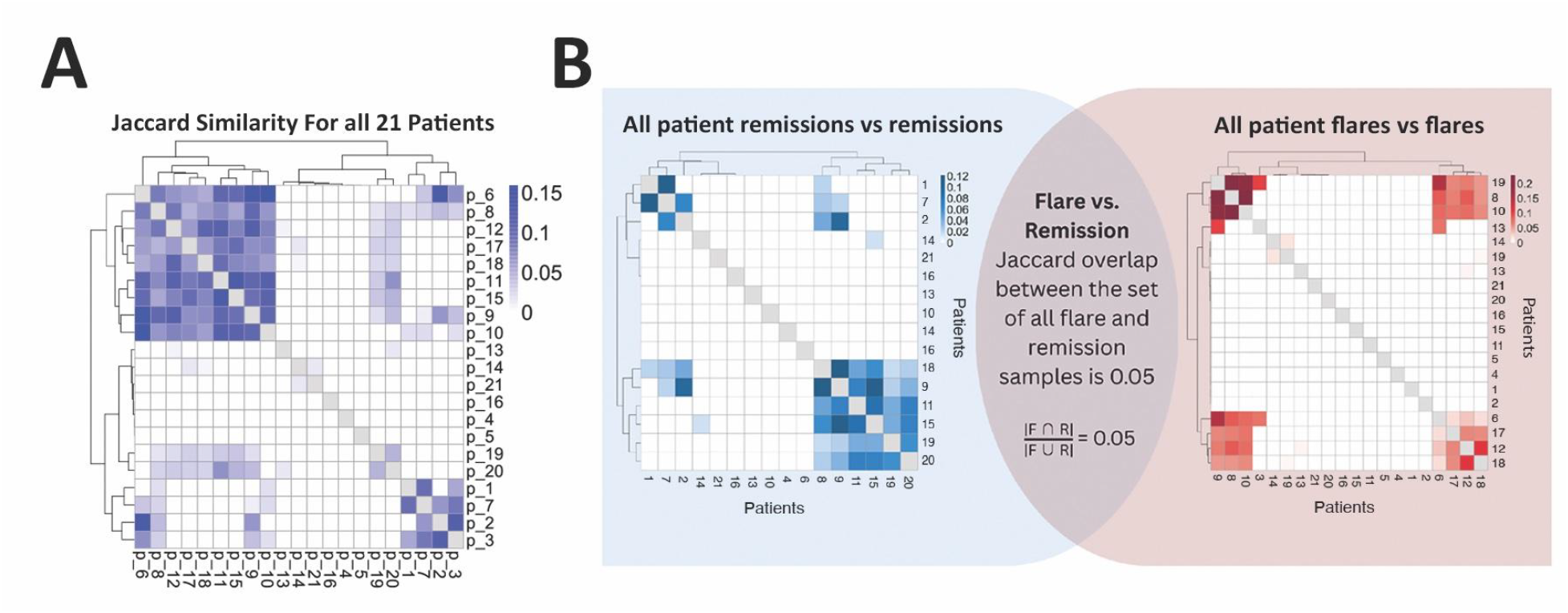
Enriched K-mers show overlaps and potential subgroups among patients. GLIPH2 was used to identify (A) enriched K-mers in all time points and (B) enriched K-mers in flare and remission states. Overlaps in K-mers were visualized using Jaccard scores. **Alt text:** Heatmaps showing Jaccard overlap scores for enriched K-mers in patients’ samples, as well as across flare and remission states. Subfigures are labeled A and B.

To further characterize these overlaps, enriched K-mers shared by at least two patients were analyzed. In flare samples, 45 enriched k-mers were shared across patients, 26 of which also appeared in remission states. These shared motifs may represent public clones, while the remaining 19 were unique to the flare state. Similarly, 35 enriched K-mers were shared across remission samples, with 9 being unique to remissions. These findings highlight the presence of both shared and state-specific motifs across patients.

Together, these findings demonstrate that some enriched K-mers are broadly shared across patients, potentially reflecting shared T cell responses across patients, and a substantial proportion appear to be specific to the disease state.

## DISCUSSION

This study highlights the potential for bulk sequencing of TCR□ sequences to uncover key insights regarding the immune landscape of UC. Quantifying repertoires’ clonality, diversity, and overlap revealed intrapatient patterns in TCR repertoires. However, due to significant patient heterogeneity and nuances in immune responses across individuals with the same diagnosis, these trends were insufficient to delineate disease states across our patient cohort. This highlighted the need for more advanced, subsequence level analyses to reveal underlying cohort-level trends. To this end, quantifying Vβ gene usage revealed differential gene frequencies between flare and remission states, where biased gene usage may reflect clonal expansion in response to specific antigens. Interestingly, many of the differentially expressed genes have been implicated in other immune-mediated diseases. TRBV20-1, one of the most frequently utilized TRBV genes, has been found to exhibit biased usage in a subset of rheumatoid arthritis patients and in a subset of CD4+ T cells reactive against the HLA-DQ2.5-glia-α1a peptide-HLA complex in patients with celiac disease; TRBV28 was identified in TCRs that recognize a superantigen linked to atopic dermatitis; and TRBV27 was also found to have a biased usage in ankylosing spondylitis patients’ TCR clones.^25-28^ Going forward, further research into Vβ gene usage in larger cohorts of UC patients, along with studies linking specific Vβ genes to distinct antigen types, will enhance our understanding of biased gene usage in UC and clarify its clinical significance.

Finally, enriched CDR3 K-mers were identified within particular patients. The notable increase in Jaccard similarity scores across certain repertoires when comparing enriched K-mers, as opposed to full TCR sequences, reinforces that although public clones may be limited, conserved motifs in the CDR3 region exist across patients and may reflect shared antigen specificities. The minimal overlap in enriched K-mers across flare and remission states especially suggests that clonal selection and antigen specificities could be distinct in patients’ inflamed and uninflamed states. Moreover, hierarchical clustering of patients into distinct groups based on the GLIPH analysis indicates there may be immunological features that characterize subsets of patients, even within our relatively small cohort. This finding motivates subsequent analyses of patients’ clinical history and data to identify features that are associated with these GLIPH-generated clusters.

These findings are promising, and further studies with larger patient cohorts will be critical to validate our observations. The predictive nature of observed features (ex. specific K-mer motifs) is unclear with 21 patients, but larger studies may elucidate whether these features can be used to distinguish flare and remission TCR repertoires in general. Additionally, because biopsies could only be obtained during patients’ regular clinical care, there were varying time intervals and numbers of samples for individual patients. Collecting PBMCs and performing bulk TCR sequencing could be a less invasive and more accessible approach to collect large amounts of samples, and previous researchers have done this; nonetheless, TCR profiling of colonic tissue in this study presents a rare opportunity to survey the immune environment in the native tissue.^11,29^ Finally, HLA typing and B cell receptor sequencing could allow for improved clustering of TCRs by shared antigen specificity, as well as provide complementary information to TCR sequencing.

Importantly, our work revealed that many patients have unique trends in Vβ gene usage or particular recurrent K-mers. Though these findings cannot be extrapolated to multiple patients, they nonetheless demonstrate the potential of personalized therapies in the context of UC. Indeed, the incredible heterogeneity across samples implies that UC patients have immunologically unique diseases. In the context of cancer, such observations have led to the development of cutting-edge treatments like TCR-engineered T cell therapies, the first of which was FDA-approved in 2024 for unresectable or metastatic synovial sarcoma.^30,31^ As TCR repertoire characterization and functional studies advance, similarly personalized immunotherapies may become feasible for patients with UC.

## REFERENCES

1. Roushan N, Ebrahimi Daryani N, Azizi Z, Pournaghshband H, Niksirat A. Differentiation of Crohn’s disease and ulcerative colitis using intestinal wall thickness of the colon: A Diagnostic accuracy study of endoscopic ultrasonography. Medical Journal of the Islamic Republic of Iran. 2019;33:57. doi:10.34171/mjiri.33.57

2. Le Berre C, Honap S, et al. Ulcerative colitis. The Lancet. 2023;(402)10401:571-584. doi:10.1016/S0140-6736(23)00966-2

3. Kobayashi T, Siegmund B, Le Berre C, et al. Ulcerative colitis. Nature Review Disease Primers. 2020;(6):74. doi:10.1038/s41572-020-0205-x

4. Croft A, Lord A, Radford-Smith G. Markers of systemic inflammation in acute attacks of ulcerative colitis: what level of C-reactive protein constitutes severe colitis?. Journal of Crohn’s and Colitis. 2022;16(7):1089–1096. doi:10.1093/ecco-jcc/jjac014

5. Hong SN, Park JY, Yang SY, Lee C, Kim YH, Joung JG. Reduced diversity of intestinal T-cell receptor repertoire in patients with Crohn’s disease. Frontiers in cellular and infection microbiology. 2022;12:932373. doi:10.3389/fcimb.2022.932373

6. Park JJ, Lee KAV, Lam SZ, et al. Machine learning identifies T cell receptor repertoire signatures associated with COVID-19 severity. Communications Biology. 2023;6(1):76. doi:10.1038/s42003-023-04447-4

7. Amoriello R, Chernigovskaya M, Greiff V, et al. TCR repertoire diversity in Multiple Sclerosis: High-dimensional bioinformatics analysis of sequences from brain, cerebrospinal fluid and peripheral blood. EBioMedicine. 2021;68. doi: 10.1016/j.ebiom.2021.103429

8. Zheng M, Zhang X, Zhou Y, et al. TCR repertoire and CDR3 motif analyses depict the role of αβ T cells in Ankylosing spondylitis. EBioMedicine. 2019;47:414–426. doi: 10.1016/j.ebiom.2019.07.032

9. Valpione, S., Mundra, P.A., Galvani, E. et al. The T cell receptor repertoire of tumor infiltrating T cells is predictive and prognostic for cancer survival. Nature Communications. 2021;(12):4098. doi: 10.1038/s41467-021-24343-x

10. Werner L, Nunberg MY, Rechavi E, et al. Altered T cell receptor beta repertoire patterns in pediatric ulcerative colitis. Clinical & Experimental Immunology. 2019;196(1):1–11. doi:10.1111/cei.13247

11. Rosati E, Pogorelyy MV, Dowds CM, et al. Identification of disease-associated traits and clonotypes in the T cell receptor repertoire of monozygotic twins affected by inflammatory bowel diseases. Journal of Crohn’s and Colitis. 2020;14(6):778–790. doi:10.1093/ecco-jcc/jjz210

12. Briggs KC, Lin JS, Chaaban L, et al. Longitudinal T cell repertoire analysis reveals dynamic clonal T cell populations in Ulcerative Colitis. bioRxiv. 2025;2025.01.13.632427. doi:10.1101/2025.01.13.632427

13. Ikeya K, Hanai H, Sugimoto K, et al. The Ulcerative Colitis Endoscopic Index of Severity More Accurately Reflects Clinical Outcomes and Long-term Prognosis than the Mayo Endoscopic Score. Journal of Crohn’s and Colitis. 2016;10(3):286–295. doi:10.1093/ecco-jcc/jjv210

14. Cottrell T, Zhang J, Zhang B, et al. Evaluating T-cell cross-reactivity between tumors and immune-related adverse events with TCR sequencing: pitfalls in interpretations of functional relevance. Journal for immunotherapy of cancer. 2021;9(7):e002642. doi:10.1136/jitc-2021-002642

15. Glanville J, Huang H, Nau A, et al. Identifying specificity groups in the T cell receptor repertoire. Nature. 2017;547:94–98 doi:10.1038/nature22976

16. turboGliph. https://github.com/HetzDra/turboGliph. Accessed May 08, 2025.

17. Chao A. Nonparametric estimation of the number of classes in a population. Scandinavian Journal of Statistics. 1984;265–270. doi:10.2307/4615964

18. Shan Y, Lee M, Chang EB. The gut microbiome and inflammatory bowel diseases. Annual review of medicine. 2022;73(1):455–468. doi: 10.1146/annurev-med-042320-021020

19. Turner SJ, La Gruta NL, Kedzierska K, Thomas PG, Doherty PC. Functional implications of T cell receptor diversity. Current opinion in immunology. 2009;21(3):286–290. doi:10.1016/j.coi.2009.05.004

20. Morisita M. Measuring of interspecific association and similarity between assemblages. Memoirs Faculty of Science Kyushu University Series E Biology. 1959;3:65–80.

21. Dash P, Fiore-Gartland A, Hertz T, et al. Quantifiable predictive features define epitope-specific T cell receptor repertoires. Nature. 2017;547:89–93. doi: 10.1038/nature22383

22. Springer I, Tickotsky N, Louzoun Y. Contribution of T cell receptor alpha and beta CDR3, MHC typing, V and J genes to peptide binding prediction. Frontiers in immunology. 2021;(12):664514. doi:10.3389/fimmu.2021.664514

23. Moeckel C, Mareboina M, Konnaris M A, et al. A survey of k-mer methods and applications in bioinformatics. Computational and structural biotechnology journal. 2024. doi:10.1016/j.csbj.2024.05.025

24. Ostmeyer J, Christley S, Toby I T, et al. Biophysicochemical motifs in T-cell receptor sequences distinguish repertoires from tumor-infiltrating lymphocyte and adjacent healthy tissue. Cancer research. 2019;79(7):1671–1680. doi: 10.1158/0008-5472.CAN-18-2292

25. Turcinov S, Af Klint E, Van Schoubroeck B, et al. Diversity and Clonality of T Cell Receptor Repertoire and Antigen Specificities in Small Joints of Early Rheumatoid Arthritis. Arthritis & Rheumatology. 2023;75(5):673–684. doi:10.1002/art.42407

26. Yohannes, DA, Freitag TL, de Kauwe A, et al. Deep sequencing of blood and gut T-cell receptor β-chains reveals gluten-induced immune signatures in celiac disease. Scientific Reports. 2017;7(1):17977. doi:10.1038/s41598-017-18137-9

27. Bryan E, Teague JE, Eligul S, et al. Human skin T cells express conserved T-cell receptors that cross-react with staphylococcal superantigens and CD1a. Journal of Investigative Dermatology. 2024;144(4):833–843. doi:10.1016/j.jid.2023.09.284

28. Hanson AL, Nel HJ, Bradbury L, et al. Altered Repertoire Diversity and Disease-Associated Clonal Expansions Revealed by T Cell Receptor Immunosequencing in Ankylosing Spondylitis Patients. Arthritis & Rheumatology. 2020;72(8):1289–1302. doi:10.1002/art.41252

29. Wu J, Pendegraft AH, Byrne-Steele M, et al. Expanded TCRβ CDR3 clonotypes distinguish Crohn’s disease and ulcerative colitis patients. Mucosal Immunology. 2018;11(5):1487–1495. doi:10.1038/s41385-018-0046-z

30. Mullard, A. FDA approves first TCR-engineered T cell therapy, for rare soft-tissue cancer. Nature Reviews Drug Discovery. 2024; 23, 731. doi:10.1038/d41573-024-00134-z

31. D’Angelo SP, Araujo DM, Abdul Razak AR, et al. Afamitresgene autoleucel for advanced synovial sarcoma and myxoid round cell liposarcoma (SPEARHEAD-1): an international, open-label, phase 2 trial. The Lancet. 2024;403(10435):1460–1471. doi:10.1016/S0140-6736(24)00319-2

